# Preparing for the next Pandemic: Simulation-based Deep Reinforcement Learning to discover and test multimodal control of systemic inflammation using repurposed immunomodulatory agents

**DOI:** 10.1101/2022.07.25.501428

**Authors:** Chase Cockrell, Dale Larie, Gary An

## Abstract

**Background:** Preparation to address the critical gap in a future pandemic between non-pharmacological measures and the deployment of new drugs/vaccines requires addressing two factors: 1) finding virus/pathogen-agnostic pathophysiological targets to mitigate disease severity and 2) finding a more rational approach to repurposing existing drugs. It is increasingly recognized that acute viral disease severity is heavily driven by the immune response to the infection (“cytokine storm”). There exist numerous clinically available biologics that suppress various pro-inflammatory cytokines/mediators, but it is extremely difficult to identify clinically effective treatment regimens with these agents. We propose that this is a complex control problem that resists standard methods of developing treatment regimens and accomplishing this goal requires the application of simulation-based, model-free deep reinforcement learning (DRL) in a fashion akin to training successful game-playing artificial intelligences (AIs). This proof-of-concept study determines if simulated sepsis (e.g. infection-driven cytokine storm) can be controlled in the absence of effective antimicrobial agents by targeting cytokines for which FDA-approved biologics currently exist.

**Methods:** We use a previously validated agent-based model, the Innate Immune Response Agent-based Model (IIRABM), for control discovery using DRL. DRL training used a Deep Deterministic Policy Gradient (DDPG) approach with a clinically plausible control interval of 6 hours with manipulation of six cytokines for which there are existing drugs: Tumor Necrosis Factor (TNF), Interleukin-1 (IL-1), Interleukin-4 (IL-4), Interleukin-8 (IL-8), Interleukin-12 (IL-12) and Interferon-*γ* (IFNg).

**Results:** DRL trained an AI policy that could improve outcomes from a baseline mortality rate of 41% (= recovery rate of 59%) to one with a recovery rate of 82.3% over 42 days simulated time.

**Discussion:** The current proof-of-concept study demonstrates that significant disease severity mitigation can potentially be accomplished with existing anti-mediator drugs, but only through a multi-modal, adaptive treatment policy requiring implementation with an AI. While the actual clinical implementation of this approach is a projection for the future, the current goal of this work is to inspire the development of a research ecosystem that marries what is needed to improve the simulation models with the development of the sensing/assay technologies to collect the data needed to iteratively refine those models.

## Introduction

A striking feature of the COVID-19 pandemic in its early phases was that medical resources, particularly those in critical care units, were overwhelmed. This issue arose primarily because of the inability to affect the underlying biological processes that drove the course of disease; once the disease manifested the only option was supportive care until the disease ran its course. Given the challenges in developing specific antiviral agents and the mandatory time required to bring novel drugs or new vaccines to clinical deployment, preparation for the next pandemic should include the development of measures that can more effectively and efficiently use existing drugs to reduce and mitigate disease severity. Specifically, with respect to COVID-19, there was an early recognition that severe disease was associated with “cytokine storm” (1-6), namely that the body’s inflammatory/immune response was producing unintended and detrimental collateral damage in response to the viral infection. There is a suggestion that based on the pathophysiological time courses of acute viral infections (1-6) disease manifestation occurs subsequent to the peak(s) of viremia; it is exactly this host-response driven pathophysiological phase that significantly contributes to in-hospital and critical care resources. As a result, there was a great deal of interest in repurposing immunomodulatory agents to attempt to mitigate disease severity (7-9), but to date, with the exception of the use of steroids for severe disease (10), none of these approaches have been unambiguously proven to be effective.

This should not come as a surprise. The phenomenon of collateral tissue damage arising from dysregulated inflammation described as “cytokine storm” is exactly the process that drives disease severity and multiple organ failure in bacterial sepsis, for which no immunomodulatory interventions have been shown to be reliably effective (11). In fact, the current set of immunotherapies for chronic inflammatory diseases, exactly those proposed for repurposed use in COVID, were themselves repurposed from agents that initially failed in sepsis trials. We have previously reported on the challenges present in attempting to control sepsis using anti-cytokine/anti-mediator therapies, primarily stemming from the failures to recognize the dynamic complexity of the mechanistic processes ostensibly being targeted (12) and that in order to be effective the treatment of sepsis should be considered a complex control problem (13). In previous work we have shown that sepsis is potentially controllable by discovering multi-modal control strategies using different types of machine learning (ML) methods trained on a complex agent-based model of acute systemic inflammation (the Innate Immune Response Agent-based Model, or IIRABM (14)) (15-17). Specifically, the latter projects described in Refs (16, 17) utilized the method, Deep Reinforcement Learning (DRL), employed by ML/Artificial Intelligence (AI) systems to successfully play and win a series of games against human experts (18-22). We term this approach *simulation-based model-free DRL*, and in prior work applied to method where we treated the attempt to control sepsis as a “game” to be played using the IIRABM, where potential cytokine interventions represented the “moves” implemented by the AI agent (16, 17).

In the context of developing capabilities to mitigate the disruption of a future potential pandemic, we make the following assertions:

1. Dysregulated and detrimental systemic inflammation is a primary source of disease severity in acute viral illness;
2. There is a critical need to have virus-agnostic disease mitigation therapies in the early phases of a pandemic; and
3. There is proven inefficacy of standard approaches to applying immunomodulation in the face of cytokine storm/sepsis; and
4. There is a need to increase the efficiency and efficacy of repurposing existing immunomodulatory agents in order to provide virus-agnostic disease mitigation options.

We propose that simulation-based control discovery using DRL can provide useful insights and potentially critical capabilities in designing effective multi-modal and adaptive immunomodulatory therapies for infections for which no effective anti-microbial agents exist. We have previously demonstrated in a proof-of-concept report that such a control policy can be discovered with DRL when manipulating up to 11 different mediators and soluble factors every 6 minutes (17). We now extend that study to evaluate whether DRL can train an artificial neural network (ANN) to discover a treatment policy utilizing existing anti-cytokine drugs to improve the outcomes to simulated infection in the absence of anti-microbial treatment.

## Methods

### Description of IIRABM

The simulation model used for DRL ANN training is a previously validated agent-based model of sepsis, the Innate Immune Response agent-based model (IIRABM) (12, 14). We have previously used the IIRABM as a surrogate/proxy system for the investigation of potential control strategies (23) for sepsis, both using genetic algorithms (15) and DRL (16, 17). A detailed description of the IIRABM can be found in Ref (14); here we present a brief overview to provide enough background to describe the control discovery work that is the subject of this paper.

The IIRABM is a two-dimensional abstract representation of the human endothelial-blood interface with the modeling assumption that the endothelial-blood interface is the initiation site for acute inflammation. The closed nature of the circulatory surface can be represented as a torus, and the two-dimensional surface of the IIRABM therefore represents the sum-total of the capillary beds in the body. The spatial scale of the real-world system is not directly mapped using this scheme. The IIRABM simulates the cellular inflammatory signaling network response to injury/infection and reproduces all the overall clinical trajectories of sepsis (14). The IIRABM incorporates multiple cell types and their interactions: endothelial cells, macrophages, neutrophils, TH0, TH1, and TH2 cells as well as their associated precursor immune cells. A schematic of the components and interactions in the IIRABM can be seen in Figure 1.

**Figure 1.**
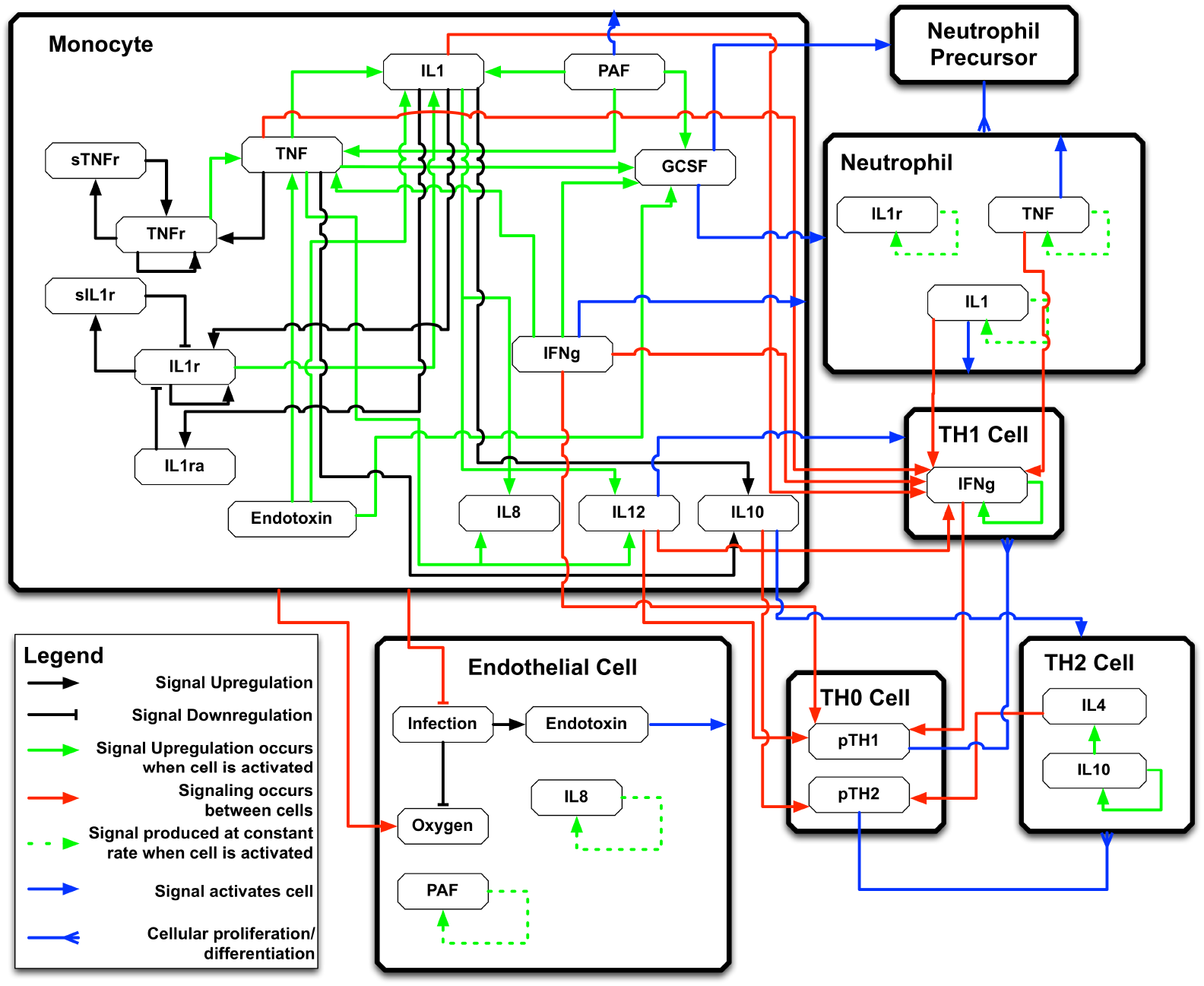
Schematic of cell types, mediator and connections in the Innate Immune Response Agent-based Model (IIRABM). IL1 = Interleukin-1, IL4 = Interleukin-4, IL8 = Interleukin-8, IL10 = Interleukin-10, IL12 = Interleukin-12, TNF = Tumor Necrosis Factor, GCSF = Granulocyte Colony Stimulation Factor, IFNg = Interferon-gamma, TNFr = Tumor necrosis factor receptor, sTNFr = soluble Tumor Necrosis Factor Receptor, IL1r = Interleukin-1 receptor, IL1ra = Interleukin-1 receptor antagonist, pTH1 = pro-Type 1 Helper T-cell state, pTH2 = pro-Type 2 Helper T-Cell state. For full details of the IIRABM see Ref (14). Figure reprinted from Ref (15) under the Creative Commons License.

The content of the IIRABM is not intended to be a comprehensive list of all the cellular subtypes present in the immune system, but rather represents the minimally sufficient set of cell populations able to represent every necessary function in the innate response to infection. System mortality of the IIRABM is defined when the aggregate endothelial cell damage, represented by the model variable “Oxy-deficit”) exceeds 80% of the total baseline health of the system; this threshold represents the ability of current medical technologies to keep patients alive (i.e., through organ support machines) in conditions that previously would have been lethal. Infectious insults to the IIRABM are initiated using 5 parameters representing the size and nature of the injury/infection as well as a metric of the host’s resilience: initial injury size, microbial invasiveness, microbial toxigenesis, environmental toxicity, and host resilience. Previous work (12) identified the boundary conditions for these parameters in terms of generating clinically realistic behavior, and therefore we consider this parameter space as representing the clinically plausible space of human response to infection.

### Deep Reinforcement Learning

Deep Deterministic Policy Gradient (DDPG) (24) was used to discover a controls algorithm that is able to heal *in silico* patients by either augmenting or diminishing the concentration of cytokine signaling molecules in the simulation. DDPG is a powerful reinforcement learning (RL) algorithm able to use off-policy data and the Bellman equation (Equation 1) to learn a value function, or Q-function, to determine the most valuable action to take given any state of the simulation.

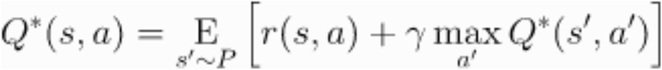

Equation 1: The Bellman Equation. Value *Q* is a function of the current state and action *(s, a)*, and is equal to the reward *r* from the current state and chosen action *(s, a)* summed with the discounted value of the next state (discount factor = γ) and action *(s’, a’)* where the next state is sampled from a probability distribution *(s’ ∼ P)*.

The Q-function is discovered through trial and error and allowing an RL agent to optimize the Q-function based on observed rewards from chosen actions. DDPG can be thought of as an extension of the Q-learning algorithm (25), where it is able to choose from a continuous action space. In Q-learning, the next action is chosen from a set of discrete actions that can be taken based on the output of the Q-function. The best action to take from the current state is identified by finding which action will return the highest value from the Q-function. Q-learning is an off-policy algorithm, which means that in the training phase, the RL agent is sometimes able to choose actions that are not the ones chosen by the Q-function. This allows the agent to explore and potentially discover actions that can lead to a greater reward than continuing from an already discovered policy. Q-learning has proven to be very powerful at solving control problems in discrete space and has proven on benchmark RL problems that the discovered controls algorithm can be very robust (24).

DDPG extends Q-learning to a continuous action space. It is too computationally expensive to exhaustively search the action space for the optimal action during the learning phase since the action space is continuous. Because of this, DDPG uses an “actor” neural network to choose an action based on the current state. The chosen action is used for the simulation, and a new state and reward is returned by the RL environment. The reward then updates the Q-function to more closely approximate the true value function for the environment, and the updated Q-function is used to perform gradient descent on the actor network to improve its decision making in the future. Because updates to the actor network are made based on an approximation, DDPG is sometimes susceptible to variations in starting conditions, and is sometimes unstable as it learns. Because of this, learning rates for the actor network and the Q-function approximation are usually slow. Additionally, to help stability, DDPG uses what is called an “Experience Replay Buffer” to sample a batch of states and actions the agent has taken in the past, instead of relying only on the current state and action for network updates.

### Training Environment

The goal of this work is to determine if an effective immunomodulatory strategy utilizing a set of existing anti-cytokine drugs could balance the need for an effective immune response to contain an infection in the absence of anti-microbials while preventing system death due to cytokine storm. We wanted to mimic a clinically relevant population and therefore chose parameter values and initial conditions to provide an overall mortality of ∼40%. Using our previously identified method of finding relevant parameter sets within bioplausible parameter space (12) we chose the following External parameters and initial infection level in order to identify the baseline conditions of the IIRABM that would subsequently be used for DRL:

- Host Resilience [oxyheal] = 0.08: This represents the rate at which the baseline endothelial cells recover their oxy level, back to a baseline of 100.
- Invasiveness [infectSpread] = 2: This represents the number of adjacent grid spaces the infection spreads to after it has reaches the carrying capacity on an individual grid.
- Environmental Toxicity [numRecurInj] = 0: This represents the number of grid spaces are randomly reinfected every 24 hours, reflecting environmental contamination. In these set of simulation experiments this function was not included.
- Toxigenesis [numInfectRep] = 2: This represents the amount of damage produced by a microbe on the grid space it occupies.
- Initial Infection Amount [inj_number] = 20: This represents the radius in number of grid spaces of a circular inoculation of the infection

With these parameters the IIRABM had a mortality of 41% (59% Recovered) at the end of simulated 42 days.

### Initial and Termination Conditions

A training episode begins 12 hours after the application of the initial infection; this is to reflect the minimal necessary incubation time between exposure and initiation of any treatment. The episode ends when either the simulated patient completely heals, dies, or 10,000 time steps (= 42 days simulated time) if neither of those conditions is met.

### Observation Space

The IIRABM states exists over a discrete, 2-dimensional 101 × 101 grid. The IIRABM includes 9 cytokines, 2 soluble cytokine receptors (essentially inhibitors of their respective cytokines), population levels of 5 different cell types. The IIRABM also reports the total amount of infection in the system and the total amount of damage present in the system (as reflected by the variable “Oxy-deficit”; but for purposes of this paper this term will be called “Total System Damage” for enhanced clarity). Since the IIRABM utilizes an abstract spatial representation, the individual discrete grids are not directly translatable to any potential spatial measurement. The aggregated system levels are considered equivalent to values potentially sampled in the blood, and therefore represent the accessible information for any potential sensor or lab assay. As this work attempts to approximate what might eventually be available clinically, we assume that any circulating cytokine/soluble receptor can be measured and returned every 6 hours: this gives the system state as reflected in 11-dimensions (e.g. 9 cytokines + 2 soluble receptors represented in the IIRABM, hereafter termed “mediators”). Alternatively, since in the clinically setting there is a distinction between infectious particles in the tissue and those that spill over into the blood, we do not consider the total system infection a clinically feasible observable and this value is not included in the observation space for the DRL. Similarly, since the total amount of damage in the system is not actually a quantifiable or observable metric in the clinical patient, this value is not included in the observations used to train the DRL; this is in contrast to our prior use of DRL trained on the IIRABM (16). As such, the current DRL agent is being trained on partially-observable states of the IIRABM.

### Action space

The actions taken by the DRL agent can either be augmentation or inhibition of six mediators present in the IIRABM for which there are existing FDA-approved pharmacological agents: Tumor Necrosis Factor (TNF), Interleukin-1 (IL-1), Interleukin-2 (IL-2), Interleukin-4 (IL-4), Interleukin-8 (IL-8), Interleukin-12 (IL-12) and Interferon-*γ* (IFNg). To reflect a clinically plausible time frame in which an blood mediatory assay would be run and used to inform the administration of a drug/set of drugs, for the DRL action space interval any or all of these mediators could be manipulated every 6 hours, As a simplifying approximation of clinical pharmacological effect the duration of the effect of each intervention was simulated to last for 6 hours. An augmentation action takes the form of the addition of a continuous value from 1 to 10 to the value of a particular mediator. Inhibition takes the form of the multiplication of the existing mediator value by 0.001 to 1; this approach is done to avoid negative (or exploding, in the case of pathway augmentation) values and is consistent with the dynamics of mediator inhibition. These are reflected in the code thusly:

if action_mag > 0, action = (action_mag*9) +1 ⇒ add mediator between 1 and 10

If action_mag <0, action = action_mag + 1.001 ⇒ multiply mediator between .001 and 1

The ability to manipulate any combination of mediators present is meant to simulate the potential use of combinations of interventions, which our prior work has suggested is necessary to effectively control sepsis (15-17); the DRL approach is intended to assist in addressing the exponential combinatorial issues associated with multi-drug therapy and the additional challenge needing to modify a particular treatment application to account for the temporal heterogeneity among individuals with regards to their disease trajectories.

### Reward function

The current DRL strategy includes two types of reward functions. The first are *terminal rewards*: these are evaluated at the end of an episode and are analogous to either winning or losing the game. The current work has a positive terminal reward if the system heals: *r* = 0.999^*step*^ − 1000. whereas the negative terminal rewards if the system dies is: *r* = 0.999^*step*^ − 1000. The incorporation of the step at which the terminating condition is met is intended to reward quicker healing, penalize faster death, and not penalize prolongation of life (albeit in a diseased state). The current work also includes *intermediate or step-wise rewards*; these are reinforcing conditions to aid in learning during the course of the episode run. The intermediate reward function is:

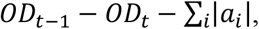

Where *OD*_*t*_ indicates the total system damage at time *t*, and *a*_*i*_ is the value for the action taken on mediator *i*. The intermediate reward calculation rewards systems that reduce their damage per time step and are able to do so with a minimal amount of intervention. The latter goal is consistent with the concept of minimizing necessary interventions and avoiding potential side effects that may not be reflected in the resolution of the simulation. Note that while we do not include the total system damage as an observable that can be used to determine actions, we believe it is valid to include it in terms of the reward function since this is a property of the simulation-based training and not intended to reflect a clinically-accessible metric. The code for the DRL environment (which includes the IIRABM and the DRL training code) can be found at https://github.com/An-Cockrell/DRL_Control

## Results

There were three possible outcomes from the simulations under control conditions: 1) Complete healing, where the Total System Damage goes to 0, 2) System Death, where the Total System Damage reaches 80% of baseline health (arbitrarily set and consistently used since initial paper on the IIRABM in 2004 (14)), and 3) “Time Out” where the system achieves neither condition #1 or #2. We note that the Time Out condition is not present in the baseline uncontrolled case, where simulated individuals either Recovered (59%) or Died (41%) by the end of the simulation run (10,000 steps = 42 days). The Time Out result arises from the application of the control policy to the simulation such that the control provides “life support” that prolongs the duration of a system run that would likely otherwise “die”. We identified that below a threshold of Total System Damage of 600 the system would invariably heal, and therefore aggregated our outcomes into two groups; “Recovered” which includes completely healed and those with Total System Damage < 600 at the end of 42 days simulated time, and “Non-recovered” for those simulations that met Death criterial (Total System Damage > 80%) or Total System Damage > 600 at the end of 42 days. Incidentally, this baseline mortality rate is approximately that of COVID-19 in the pre-pharmacological treatment era.

DRL training proceeded for 400 episodes and converged to a policy that had a Post-Control Recovered rate = 82.3% with a Non-recovered rate of 17.7% ; n = 1000 with 823 Recovered (810 Complete Recovery and 13 Timeouts with Total System Damage < 600) and 177 Non-recovered (169 Timeouts with Total System Damage > 600 and 8 Deaths). These results were a significant improvement over the uncontrolled base condition, which had a Mortality of 41% (= 59% Complete Healing/Recovered). The improvement could be considered even more significant if only “true” Deaths were counted (“true” mortality rate = 0.8%), but as noted above we recognize that a significant proportion of the Time Outs would go on to “die” if the control was stopped and the simulations continued; this can be seen in the plot of Total System Damage trajectories for both Recovered and Timeout/Death system runs in Figure 2A. There appears to be a “point of no return” if the Total System Damage exceeds ∼ 4000, at which point no control is able to steer the system back to complete recovery. Panel 2B and 2C show the total level of cytokines/mediators present in the system in both the Recovered (Figure 2B) and Non-Recovered (Figure 2C) groups. It is evident that worse outcome was associated with sustained levels of pathway activation, which intuitive corresponds to “cytokine storm.”

**Figure 2:**
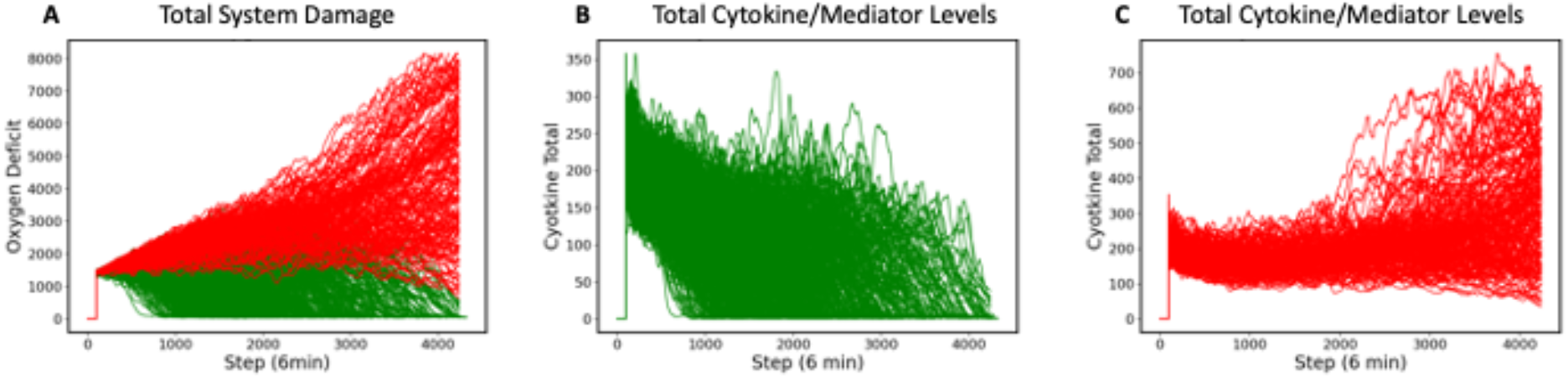
Differences between Recovered and Non-recovered Groups in terms of Total System Damage and Levels of Total Cytokines/Mediators present in the system. Panel A: Oxy-Deficit (= Total System Damage) Trajectories with DRL control policy. N = 1000, Recovered (Green) = 823 (810 Completely Healed + 13 Time-Out with Total System Damage < 600), Non-Recovered (Red) = 177 (169 Time-Out with Total System Damage > 600 and 8 True Deaths). Panel B: Trajectories of Total Cytokines/Mediators under control policy for the Recovered Group (N = 823). Panel C: Trajectories of Total Cytokines/Mediators under control policy for the Non-recovered Group (N = 173). Comparing Panels B and C it is evident that worse outcome was associated with sustained activation of the system’s pathways, lending intuitive creedance to policies that focus on mediator inhibition.

The control policy was also able to completely eradicate the initial infection without the aid of antimicrobials by augmenting the immune clearance capability (Figure 3). Note that the infection is essentially eradicated by time step ∼ 2500.

**Figure 3.**
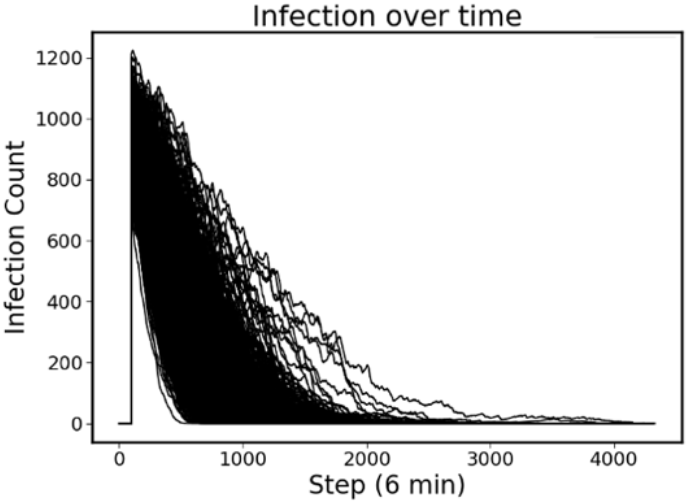
Clearance of initial infection via controlled immune functions. In all circumstances the initial infection was controlled by modulating the 6 targeted mediators, with infection essentially eradicated by Time Step ∼2500.

The discovered policy manipulated each of the 6 targeted mediators in some fashion (either augmentation or inhibition) every time step in a variable fashion. Successful control outcomes in the Recovered group (Figure 4A-F) can be grouped into those where the action was always some degree of inhibition: IL8 (Fig 4A) and IL12 (Fig 4B) versus those where some degree of augmentation was utilized at certain points: TNF (Fig 4C), IL1 (Fig 4D), IL4 (Fig 4E) and IFNg (Fig 4F). In this later group, TNF and IL1 were only augmented after infection was eradicated, a seeming non-intuitive finding that reinforces the complex nature of the control problem (see Discussion).

**Figure 4A-F.**
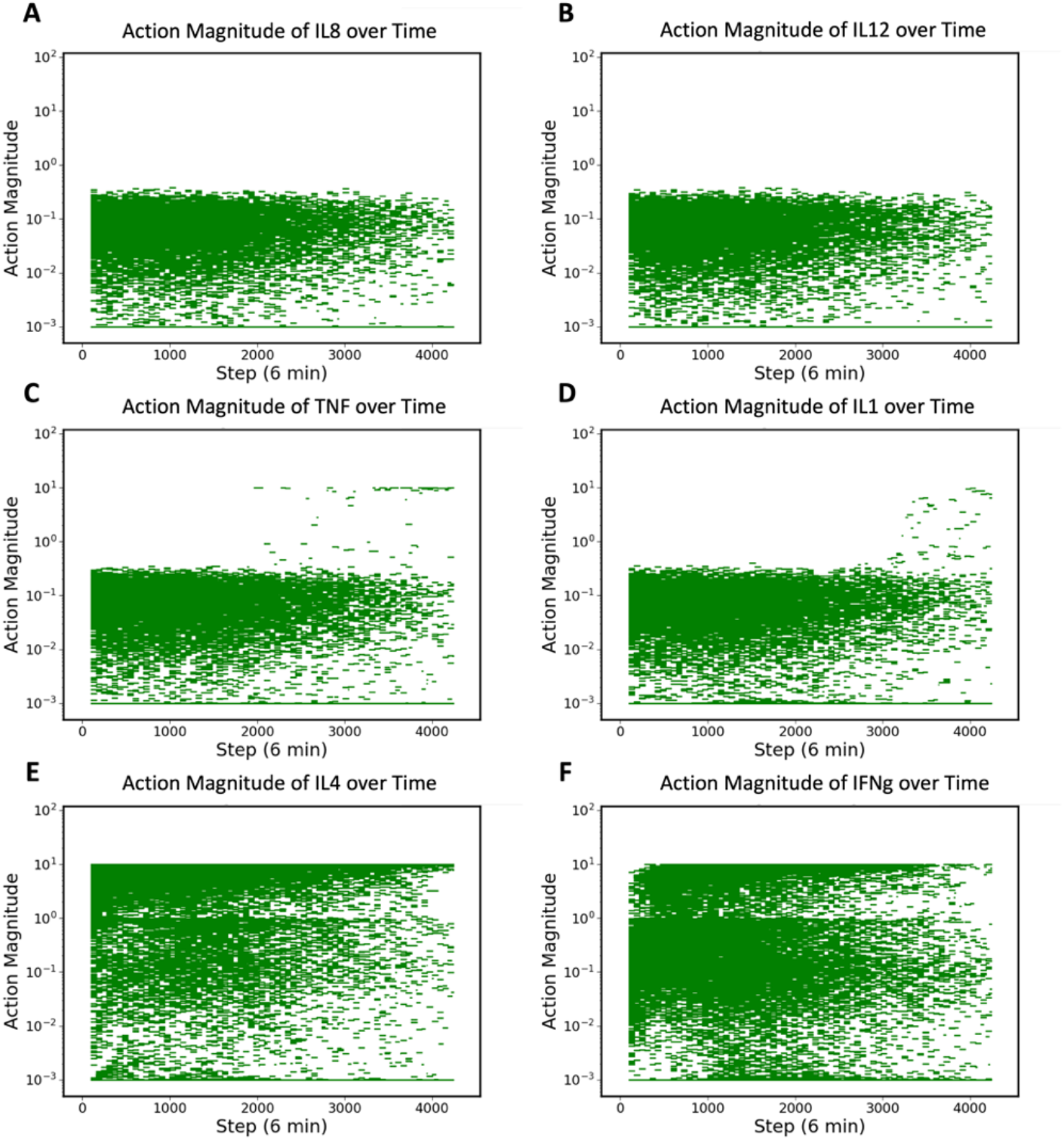
Control Actions taken in Recovery Group (N = 823). Panels A and B show control actions that always inhibit IL8 and IL12. Panels C and D show control actions that mostly inhibit TNF and IL1, with the exception of a few cases where these mediators are augmented near the end of runs. Panels E and F show control actions that can vary from augmentation to inhibition for IL4 and IFNg.

The controls applied for the Non-recovery group can be seen In Figure 5A-F; these actions have a similar pattern, with the only notable difference a greater number of extreme measures taken as the Total System Damage passes the putative point of no return (this is particularly notable in the increased circumstances of augmenting TNF and IL1),

**Figure 5A-F.**
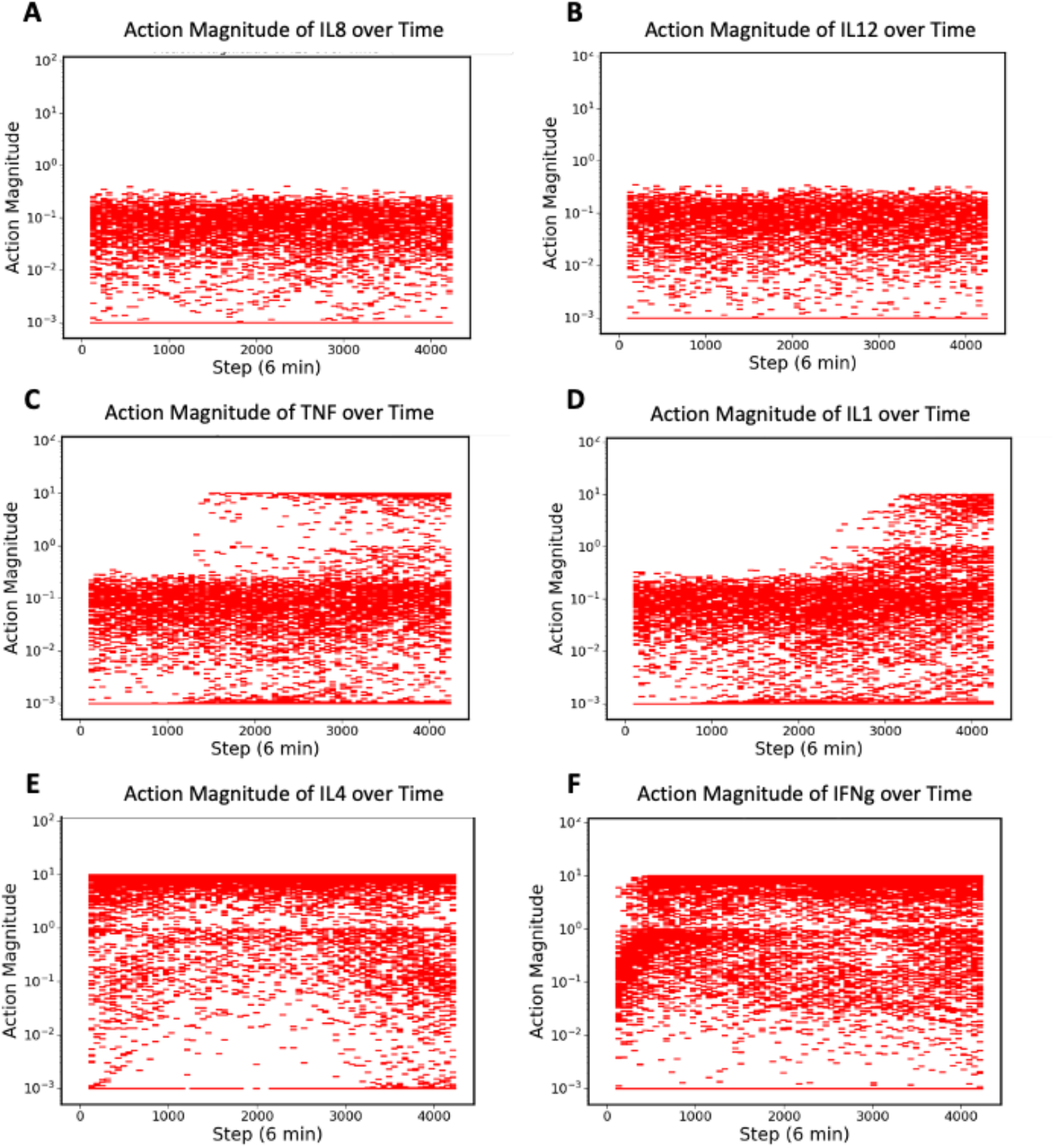
Control Actions taken in Non-recovery Group (N = 177). Panels A and B show control actions that always inhibit IL8 and IL12. Panels C and D show control actions that mostly inhibit TNF and IL1, but with an increase in the augmentation occurrences compared to Panel 4C and D for the Reocovered group. Panels E and F show control actions that can vary from augmentation to inhibition for IL4 and IFNg, also with increased augmentation compared to respective actions for IL4 and IFNg in the Recovered Group (Panels 4E and 4F).

To further visualize the effect of the various controls we compare the actions taken with the resulting effect as seen by the trajectories of the targeted mediators. Figures 6-8 shows these relationships for the Recovered group. Figure 6 shows the results of controls in the inhibition-always group of IL8 and IL12; in both these cases mediator levels are kept suppressed for the duration of the simulations.

**Figure 6.**
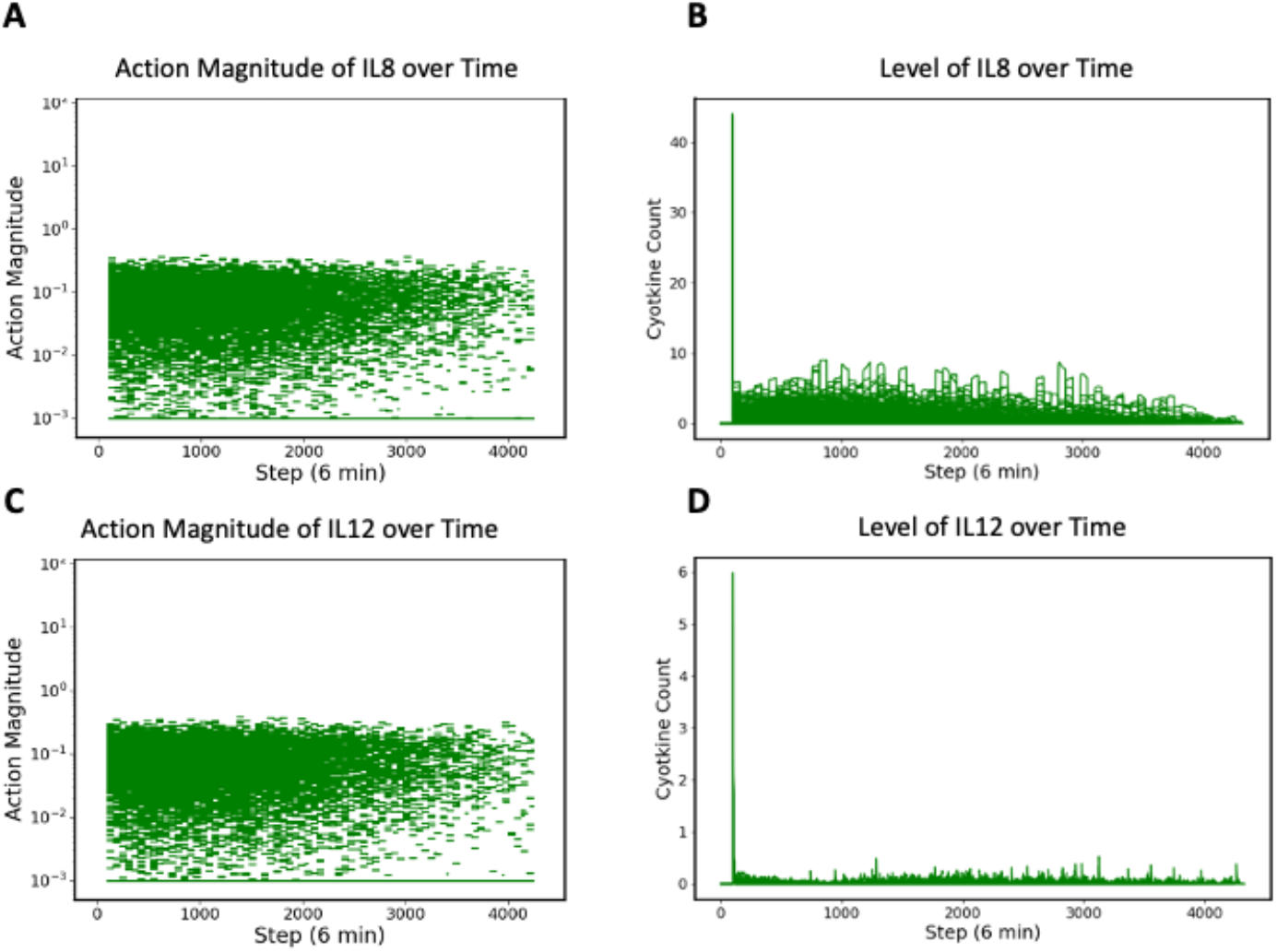
Control actions and associated mediator levels for IL8 and IL12 in the Recovered Group. Panel A shows control actions for IL8, Panel B shows levels of IL8, Panel C shows control actions for IL12 and Panel D shows levels of IL12. Control policy treats these two mediators as targets for persistent inhibition.

**Figure 7.**
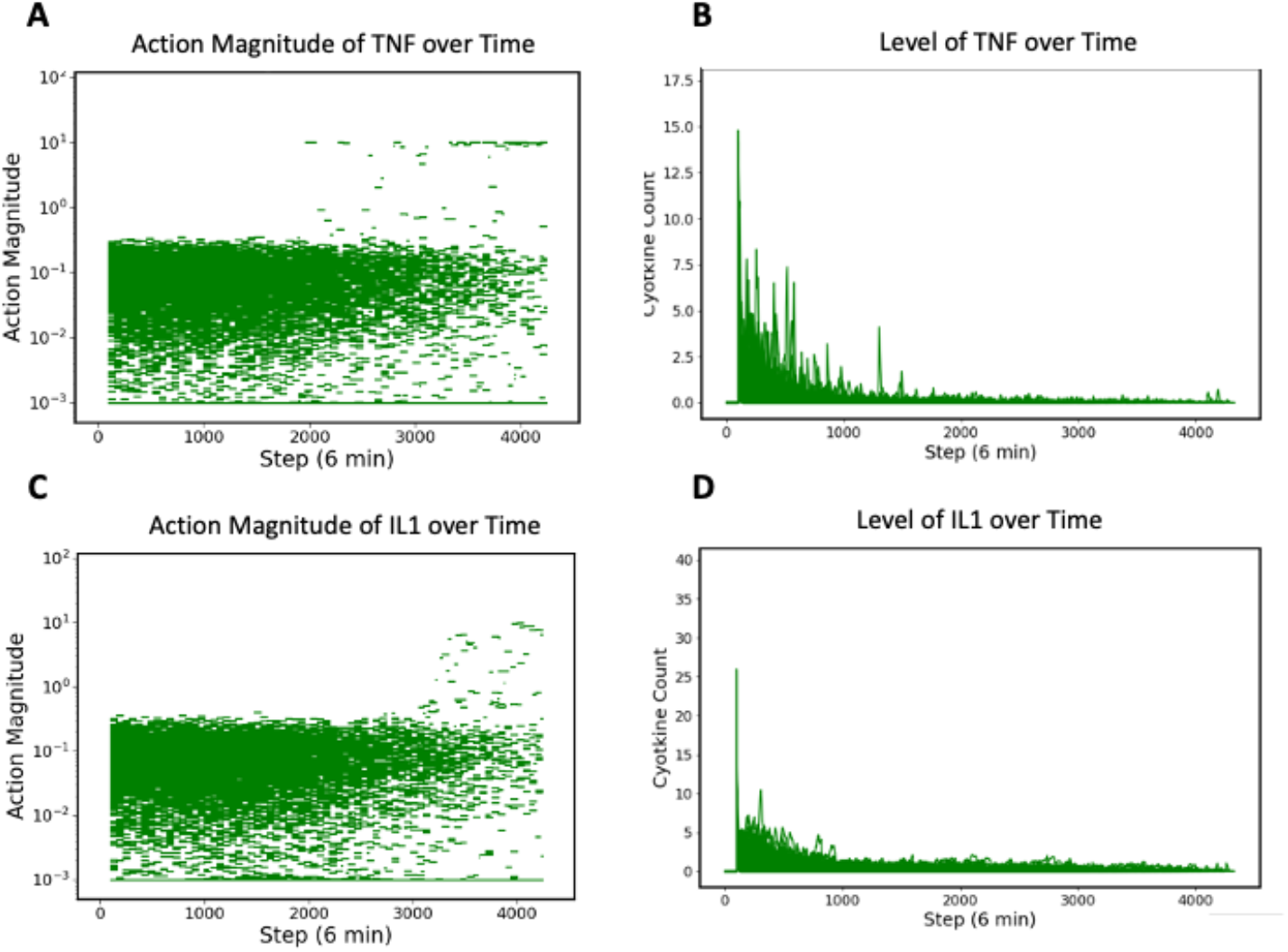
Control actions and associated mediator levels for TNF and IL1 in the Recovered Group. Panel A shows controls on TNF, Panel B shows levels of TNF, Panel C shows controls on IL1 and Panel D shows levels of IL1. Note that while the control action is primarily inhibition, there are sporadic cases of augmentation in the period after the infection has been eradicated (∼ Time Step 2500).

**Figure 8.**
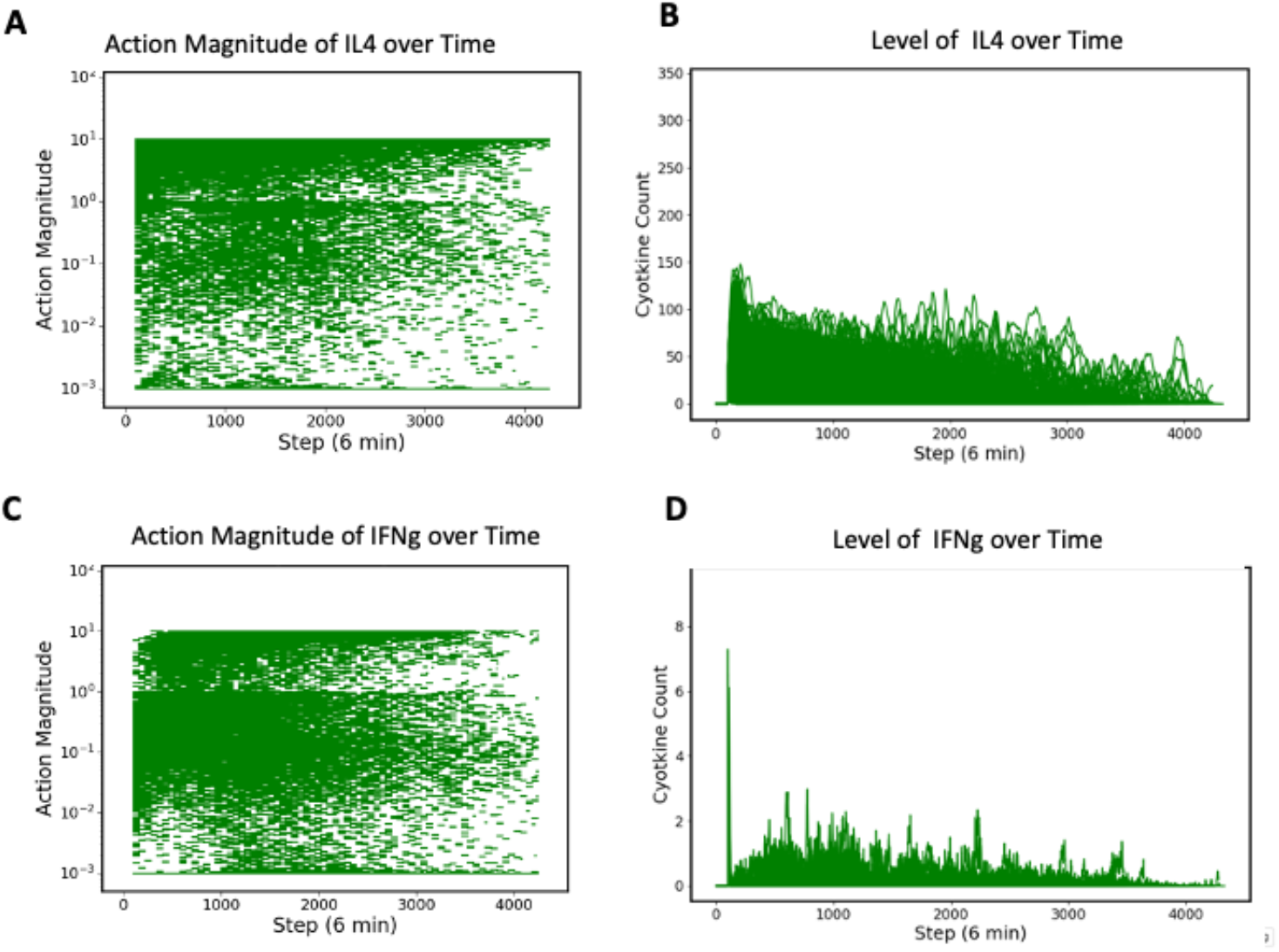
Control actions and associated mediator levels for IL4 and IFNg in the Recovered Group. Panel A shows controls on IL4, Panel B shows levels of IL4, Panel C shows controls on IFNg and Panel D shows levels of IFNg. Note these control actions range between augmentation and inhibition, with the effect of generally decreasing mediator levels over time as the system recovers.

Figure 7 shows the results of the controls on TNF and IL1, which are predominantly inhibitory with only rare instances of late supplementation. The trends show higher levels of these mediators in the early phase of infection, albeit with a steady decrease over time.

Figure 8 shows the results of controls in the variable augmentation/inhibition group of IL4 and IFNg. Both of these mediators show slightly more prolonged trajectories, but also decrease as the system heals. This is consistent with the finding that the process of healing involves the eventual resolution of mediator/cytokine activity (Figure 2B).

Figures 9-11 demonstrate the control actions and corresponding mediator trajectories in the Non-Recovery group. Examination of these can aid in the interpretation of the overall policy and how it may have been influenced by the training regimen. Figure 9 shows the control actions and trajectories for IL8 and IL12, the persistently inhibited pair of targeted mediators. This overall pattern does not look too different than the controls/trajectories in the Recovered group for IL8 and IL12, though in the later phases where the systems are moving towards the Death threshold the overall levels of these mediators are slightly greater, again consistent with the concept of non-resolving inflammatory activity as a harbinger of poor outcome.

**Figure 9.**
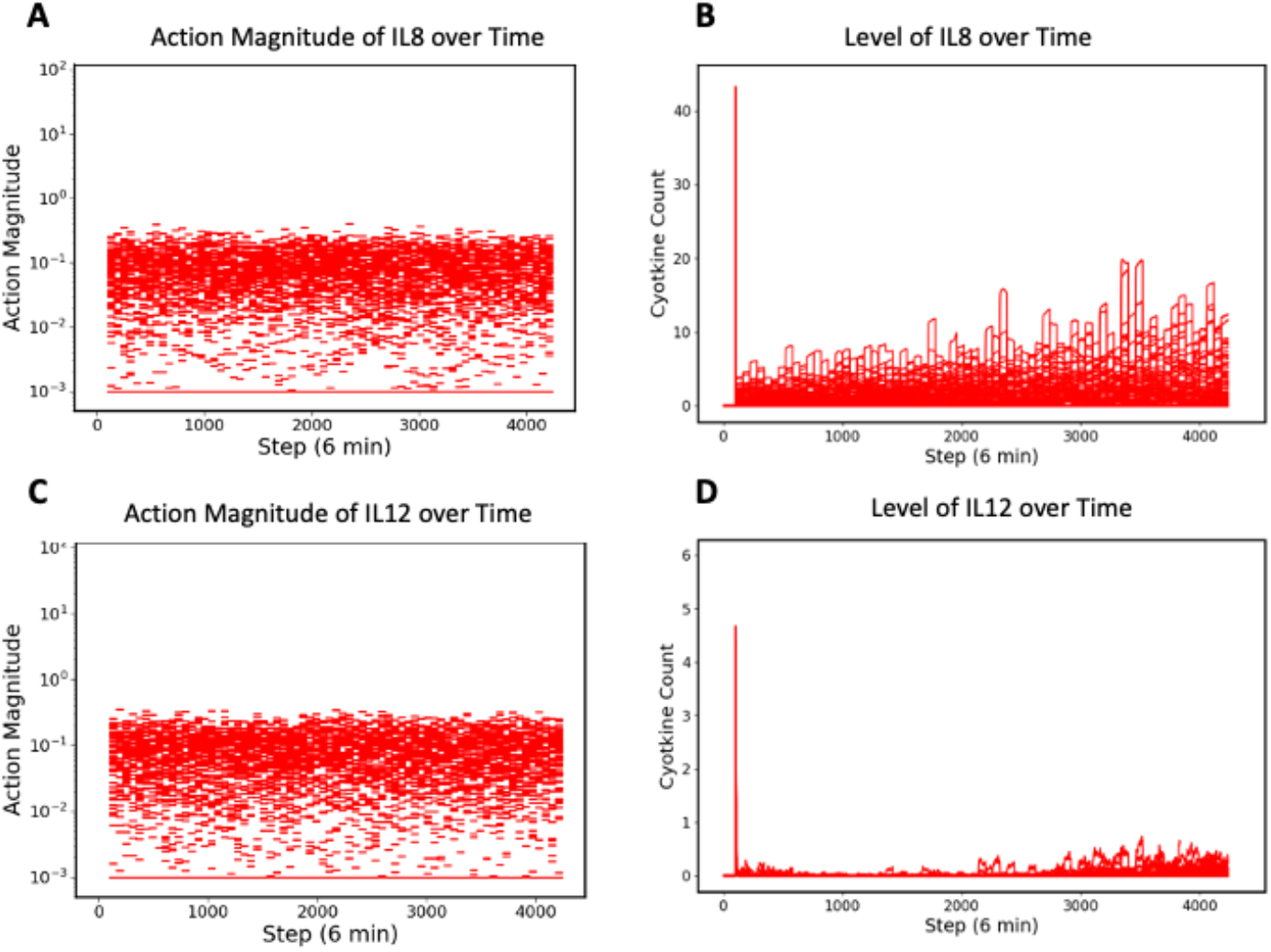
Control actions and associated mediator levels for IL8 and IL12 in the Non-Recovered Group. Panel A shows control actions on IL8, Panel B shows levels of IL8, Panel C shows control actions on IL12 and Panel D shows levels of IL12. Note that the effectiveness of inhibition appears to become less as the mediator levels slightly rise at the end of the runs. This is consistent with worsened outcomes being marked by persistent/increasing levels of pathway/mediator activity (e.g. cytokine storm).

This pattern is more pronounced in the control/trajectory plots for TNF and IL1 seen in Figure 10. In these runs the late augmentation of these two mediators is more pronounced than in the Recovered group (Figure 7). Here there is a seeming paradoxical increase in augmentation when the systems are clearly progressing towards a “cytokine storm” death. We hypothesize that what is happening here is an unintended consequence due to the overall training where earlier training episodes rewarded augmentation of pro-inflammatory mediators to eradicate the initial infection in its early phases. However, since the training observation space does not include the total infection number, the arrived-at policy cannot distinguish whether the rising mediator levels are due to worsening infection or from forward feedback cytokine storm. Ironically, the DRL finds a response policy that mimics the same paradox inherent in the biological/clinical setting: self-generated system damage from pro-inflammation.

**Figure 10.**
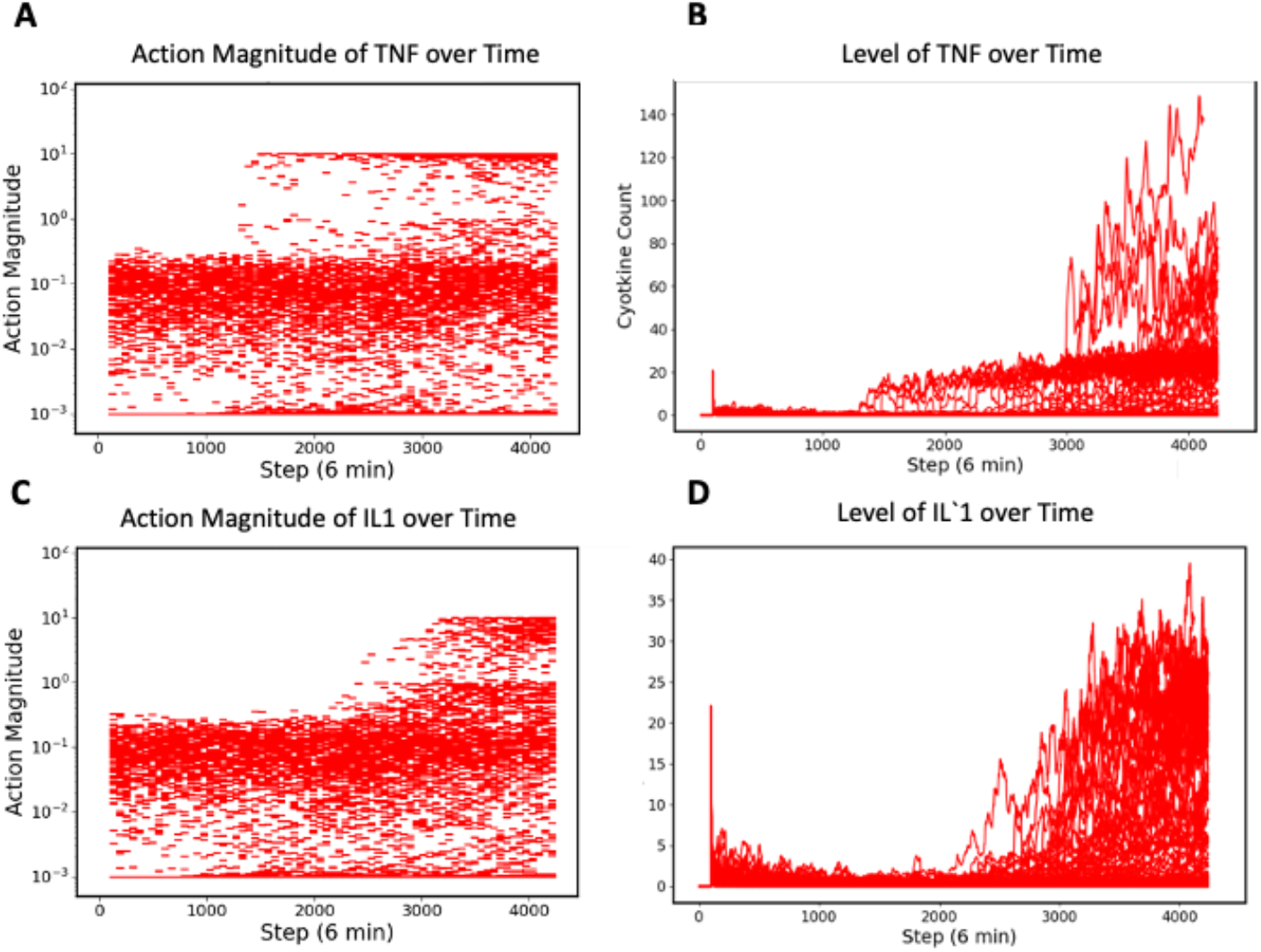
Control actions and associated mediator levels for TNF and IL1 in the Non-Recovered Group. Panel A shows control actions on TNF, Panel B shows levels of TNF, Panel C shows control actions on IL1 and Panel D shows levels of IL1. Note that there is more augmentation of these mediators at the end of the runs compared to successful controls seen in Figure 7. This is consistent with worsened outcomes being marked by persistent/increasing levels of pathway/mediator activity (e.g. cytokine storm).

A slightly different perspective of this same phenomenon can be seen in the plots of actions/trajectories of IL4 and IFNg in Figure 11. In Panels 11C and D for IFNg, which is has generally pro-inflammatory promoting functions, there is an increase in augmentation as the system moves to a higher mediator activity state. In Panels 11A and 11B, IL4, which primarily promotes the generation of anti-inflammatory TH2 T-cells, can be seen increasing both augmentation and mediator levels during the same period of time. As the sole targeted mediator in the DRL’s action space that primarily increases anti-inflammation, IL4 may be considered as the only available counter-action to the other pro-inflammatory mediators present.

**Figure 11.**
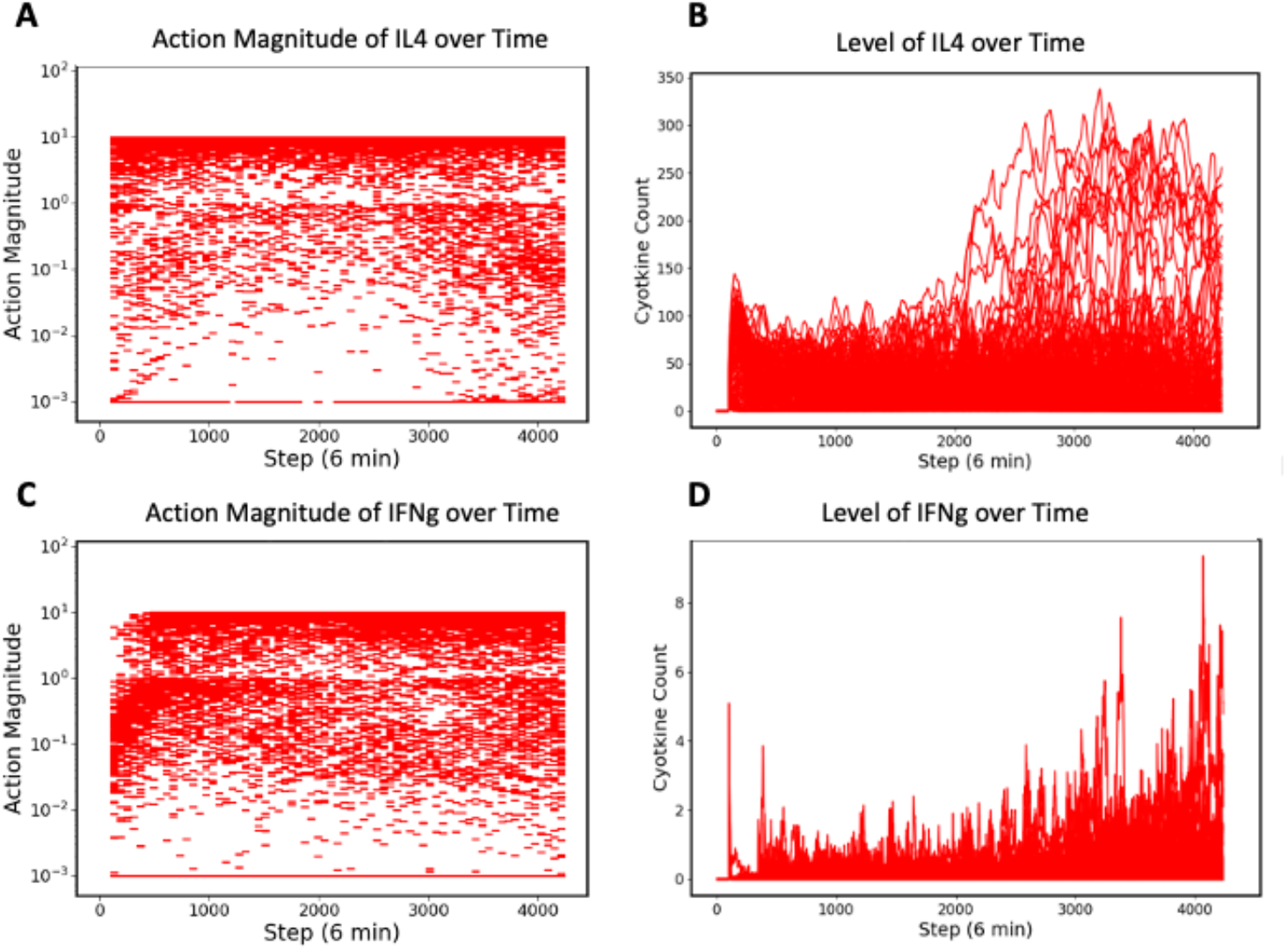
Control actions and associated mediator levels for IL4 and IFNg in the Non-Recovered Group. Panel A shows control actions on IL4, Panel B shows levels of IL4, Panel C shows control actions on IFNg and Panel D shows levels of IFNg. As with the other Non-recovered mediator trajectories, values increase as the system progress towards death. The same pattern of undesirable augmentation occurs with IFNg as with the other pro-inflammatory mediators, though within the priorly implemented action range. IL4, the sole generally anti-inflammatory mediator in the current control space, is also mostly augmented; while this could be considered appropriate it is insufficient and may suggest additional anti-inflammatory mediators would need to be targeted.

## Summary

Future pandemics are an inevitability, and while many preparations for future pandemics focus on somehow enhancing novel drug/anti-viral/vaccine development (important as they are), there are certain points that are worth noting:

1. Pathogen-agnostic disease mitigation is a critical capability in terms of readiness for future viral pandemics. While there is a certain appeal to developing viral-species specific interventions, such as anti-viral agents and vaccines, these agents have a mandatory lag-time in terms of their development; despite the impressive and unprecedented success and rapidity of COVID-19 vaccine development, it is difficult to imagine how such modalities could be made available is less than a year. Alternatively, there is a highly conserved mechanism of disease pathogenesis arising from the host inflammatory response, a shared feature of many viral infections (1-6). Developing effective strategies to control this process, while maintaining host capability to eradicate the infection, would provide a crucial capability in the early phases of any future pandemic.
2. However, the need to balance effective inflammatory/immune antimicrobial responses while mitigating the detrimental effects excessive inflammation is a highly complex task. Sepsis has been known to involve disordered and “excessive” inflammation for half a century (26). However, attempts to modulate the inflammatory response in the face of acute infection ever since have failed to effectively translate into the clinical arena (12). COVID-19 resurrected this interest (27), with what should have been expected undecisive results. The general failure of immunomodulation in the face of acute infection suggests that future approaches should consider this problem as complex control problem, and apply methods appropriate to solving complex control problems (13).
3. Drug repurposing is not as simple as extrapolating the putative mechanism of a drug and assuming that such a mechanism would be efficacious in a completely different context. The urgency of COVID-19 prompted the initiation of multiple potential therapies and trials based on bioplausibility; but it should be noted that every failed clinical trial presupposes that same bioplausibility. The same Translational Dilemma present in the development of new therapeutics (28) is also in play with the drug repurposing task, and requires the same readjustment of how to accomplish that task. Notably, the nature of the Translational Dilemma, i.e., the need to dynamically mechanistically-evaluate putative mechanistic bioplausibility, means that correlative approaches that utilize AI/traditional computational approaches do not provide a scientifically sound path that addresses the fundamental step in the drug evaluation process because they rely on correlative methods and the extrapolation of mechanistic-effect that has been demonstrated to be ineffective (29-31).

We have previously proposed that the integration of advanced forms of ML (specifically DRL) and high-fidelity mechanism-based simulations provides a scientifically sound path forward (13, 16, 32). A clear and critical challenge moving forward is developing more detailed and trustworthy simulation models that can be used for training AI-controllers that can be clinically deployed. Achieving this goal includes technical challenges that include not only being able to represent the biology in sufficient detail, but developing methods for calibrating and parameterizing such models that take into account the inherent incompleteness of biological knowledge and the considerable heterogeneity seen in biological behavior (33, 34). The need to deal with a perpetual and inevitable incompleteness of mechanistic knowledge is a key point that needs to be recognized: it cannot be that we must wait to know “everything” about how the biology works before we can hope to engineer interventions. Rather, we must recognize the need to develop paths forward that can provide some clinical utility while building in the capabilities to perpetually and iteratively improve and refine our simulation models. An example of this structure can be seen in the evolution of weather modeling and prediction: when the importance of being able to predict hurricane behavior was identified in the 1950s it was well-recognized that the existing mathematical models were insufficient for the task. But rather than saying this extremely difficult task was not pursuing, a model-driven data collection ecosystem was developed with the explicit goal of both providing some then-present-day benefit (limited though that may be), but, more importantly, identify and collect the *type of data needed to improve the models*. In this case, the modeling proposed was not limited by what sort of data could be collected; rather, the types/scale/complexity of the models needed to solve the problem were specified, and data collection strategies and capabilities were developed to allow the construction of the necessary models. This is the inflection point that biomedical community faces today in terms of fully leveraging the potential of mechanism-based, algorithmic simulation models (32, 35). We hope that the proof-of-concept demonstration presented in this manuscript will provide additional stimulus at the potential for the role of mechanistic algorithmic simulation models and how those models can be integrated with cutting edge ML and AI methods. We hope that this work will prompt additional investigations to improve and advance this methodology, and, critically, help drive the corresponding developments in real-time mediator/cytokine sensing and administration such that we will be better prepared to face the next inevitable pandemic.

## Acknowledgements

This work was supported in part by the National Institutes of Health Award UO1EB025825. This research is sponsored by the Defense Advanced Research Projects Agency (DARPA) through Cooperative Agreement D20AC00002 awarded by the U.S. Department of the Interior (DOI), Interior Business Center. The content of the information does not necessarily reflect the position or the policy of the Government, and no official endorsement should be inferred.

## References

1. de Rivero Vaccari JC, Dietrich WD, Keane RW, de Rivero Vaccari JP. The Inflammasome in Times of COVID-19. Frontiers in immunology. 2020;11:2474.

2. Hu B, Huang S, Yin L. The cytokine storm and COVID-19. Journal of medical virology. 2021;93(1):250–6.

3. Liu Q, Zhou Y-h, Yang Z-q. The cytokine storm of severe influenza and development of immunomodulatory therapy. Cellular & molecular immunology. 2016;13(1):3–10.

4. Jafarzadeh A, Chauhan P, Saha B, Jafarzadeh S, Nemati M. Contribution of monocytes and macrophages to the local tissue inflammation and cytokine storm in COVID-19: Lessons from SARS and MERS, and potential therapeutic interventions. Life sciences. 2020:118102.

5. Falasca L, Agrati C, Petrosillo N, Di Caro A, Capobianchi M, Ippolito G, et al. Molecular mechanisms of Ebola virus pathogenesis: focus on cell death. Cell Death & Differentiation. 2015;22(8):1250–9.

6. Srikiatkhachorn A, Mathew A, Rothman AL, editors. Immune-mediated cytokine storm and its role in severe dengue. Seminars in immunopathology; 2017 2017, July: Springer; 2017.

7. Ingraham NE, Lotfi-Emran S, Thielen BK, Techar K, Morris RS, Holtan SG, et al. Immunomodulation in COVID-19. The Lancet Respiratory Medicine. 2020;8(6):544–6.

8. Hertanto DM, Wiratama BS, Sutanto H, Wungu CDK. Immunomodulation as a potent COVID-19 pharmacotherapy: Past, present and future. Journal of Inflammation Research. 2021;14:3419.

9. Meyerowitz EA, Sen P, Schoenfeld SR, Neilan TG, Frigault MJ, Stone JH, et al. Immunomodulation as Treatment for Severe Coronavirus Disease 2019: A Systematic Review of Current Modalities and Future Directions. Clin Infect Dis. 2021;72(12):e1130–e43.

10. Horby P, Lim WS, Emberson JR, Mafham M, Bell JL, Linsell L, et al. Dexamethasone in Hospitalized Patients with Covid-19. N Engl J Med. 2021;384(8):693–704.

11. Chousterman BG, Swirski FK, Weber GF, editors. Cytokine storm and sepsis disease pathogenesis. Seminars in immunopathology; 2017: Springer.

12. Cockrell C, An G. Sepsis Reconsidered: Identifying Novel Metrics For Behavioral Landscape Characterization With A High-Performance Computing Implementation Of An Agent-Based Model. bioRxiv. 2017:141804.

13. An G, Cockrell C, Day J. Therapeutics as Control: Model-Based Control Discovery for Sepsis. Complex Systems and Computational Biology Approaches to Acute Inflammation: Springer; 2021. p. 71–96.

14. An G. In silico experiments of existing and hypothetical cytokine-directed clinical trials using agent-based modeling. Critical care medicine. 2004;32(10):2050–60.

15. Cockrell RC, An G. Examining the controllability of sepsis using genetic algorithms on an agent-based model of systemic inflammation. PLoS Comput Biol. 2018;14(2):e1005876.

16. Petersen BK, Yang J, Grathwohl WS, Cockrell C, Santiago C, An G, et al. Deep reinforcement learning and simulation as a path toward precision medicine. Journal of Computational Biology. 2019;26(6):597–604.

17. Larie D, An G, Cockrell C. Preparing for the next COVID: Deep Reinforcement Learning trained Artificial Intelligence discovery of multi-modal immunomodulatory control of systemic inflammation in the absence of effective anti-microbials. bioRxiv. 2022.

18. Silver D, Hubert T, Schrittwieser J, Antonoglou I, Lai M, Guez A, et al. A general reinforcement learning algorithm that masters chess, shogi, and Go through self-play. Science. 2018;362(6419):1140–4.

19. Silver D, Schrittwieser J, Simonyan K, Antonoglou I, Huang A, Guez A, et al. Mastering the game of Go without human knowledge. Nature. 2017;550(7676):354–9.

20. Vinyals O, Babuschkin I, Czarnecki WM, Mathieu M, Dudzik A, Chung J, et al. Grandmaster level in StarCraft II using multi-agent reinforcement learning. Nature. 2019;575(7782):350–4.

21. Jumper J, Evans R, Pritzel A, Green T, Figurnov M, Ronneberger O, et al. Highly accurate protein structure prediction with AlphaFold. Nature. 2021;596(7873):583–9.

22. Kirkpatrick J, McMorrow B, Turban DH, Gaunt AL, Spencer JS, Matthews AG, et al. Pushing the frontiers of density functionals by solving the fractional electron problem. Science. 2021;374(6573):1385–9.

23. An G, Fitzpatrick BG, Christley S, Federico P, Kanarek A, Neilan RM, et al. Optimization and Control of Agent-Based Models in Biology: A Perspective. Bull Math Biol. 2017;79(1):63–87.

24. Lillicrap TP, Hunt JJ, Pritzel A, Heess N, Erez T, Tassa Y, et al. Continuous control with deep reinforcement learning. arXiv preprint 150902971. 2015.

25. Mnih V, Kavukcuoglu K, Silver D, Graves A, Antonoglou I, Wierstra D, et al. Playing atari with deep reinforcement learning. arXiv preprint 13125602. 2013.

26. Bone RC. The pathogenesis of sepsis. Annals of internal medicine. 1991;115(6):457–69.

27. Parvathaneni V, Gupta V. Utilizing drug repurposing against COVID-19–efficacy, limitations, and challenges. Life sciences. 2020;259:118275.

28. An G. Closing the scientific loop: bridging correlation and causality in the petaflop age. Science translational medicine. 2010;2(41):41ps34–41ps34.

29. Zhou Y, Wang F, Tang J, Nussinov R, Cheng F. Artificial intelligence in COVID-19 drug repurposing. The Lancet Digital Health. 2020;2(12):e667–e76.

30. Mohanty S, Rashid MHA, Mridul M, Mohanty C, Swayamsiddha S. Application of Artificial Intelligence in COVID-19 drug repurposing. Diabetes & Metabolic Syndrome: Clinical Research & Reviews. 2020;14(5):1027–31.

31. Galindez G, Matschinske J, Rose TD, Sadegh S, Salgado-Albarrán M, Späth J, et al. Lessons from the COVID-19 pandemic for advancing computational drug repurposing strategies. Nature Computational Science. 2021;1(1):33–41.

32. An G. Specialty Grand Challenge: What it Will Take to Cross the Valley of Death: Translational Systems Biology, “True” Precision Medicine, Medical Digital Twins, Artificial Intelligence and In Silico Clinical Trials. Frontiers in Systems Biology. 2022;2.

33. Cockrell C, Ozik J, Collier N, An G. Nested active learning for efficient model contextualization and parameterization: pathway to generating simulated populations using multi-scale computational models. Simulation. 2021;97(4):287–96.

34. Chase Cockrell GA. Utilizing the heterogeneity of clinical data for model refinement and rule discovery through the application of genetic algorithms to calibrate a high-dimensional agent-based model of systemic inflammation. Frontiers in physiology. 2021;12.

35. An G, Cockrell C. Drug Development Digital Twins for Drug Discovery, Testing and Repurposing: A Schema for Requirements and Development. Frontiers in Systems Biology. 2022;2.

